# Intranasal oxytocin mRNA-LNP can promote social behaviour and reduce pain

**DOI:** 10.64898/2026.03.18.711938

**Authors:** Lipin Loo, Kazuma Fujikake, Maria I. Bergamasco, Rebecca Carr, Riley O’Shea, Tian Y. Du, Scott B. Cohen, Febrina Sandra, Pall Thordasson, Loren J. Martin, Celesta Fong, G. Gregory Neely

## Abstract

The COVID-19 epidemic and success of mRNA-LNP vaccines demonstrated the transformative potential of mRNA therapeutics. Beyond vaccination, mRNA delivery offers a platform for transient on-demand expression of therapeutic proteins for both rare and common diseases. While delivery of therapeutics to the liver is relatively straightforward, targeted delivery of mRNA-LNPs to the central nervous system (CNS) remains a significant challenge. Here we show that intranasal mRNA-LNP delivery results in localized mRNA cargo uptake and functional expression in the respiratory and olfactory epithelium, where the encoded cargo protein is secreted and can enter the CNS. Guided by genomic data of pain-associated gene expression, we identified secreted proteins as candidate mRNA-encoded analgesics. Intranasal mRNA-LNP encoding a synthetic oxytocin transcript (*OXT*) resulted in bioactive oxytocin peptide delivery to the CNS. Functionally, intranasal *OXT* mRNA-LNP enhanced social behaviour and attenuated pain responses across multiple behavioural paradigms, without impairing motor coordination. Importantly, repeated dosing was well tolerated and intranasal mRNA-LNP did not elicit an inflammatory response or alter overall health. Together, these findings establish intranasal mRNA-LNP delivery of secreted ligands as a safe, non-invasive route to target the CNS, unlocking a new class of mRNA therapeutics for pain or other disorders of the brain.

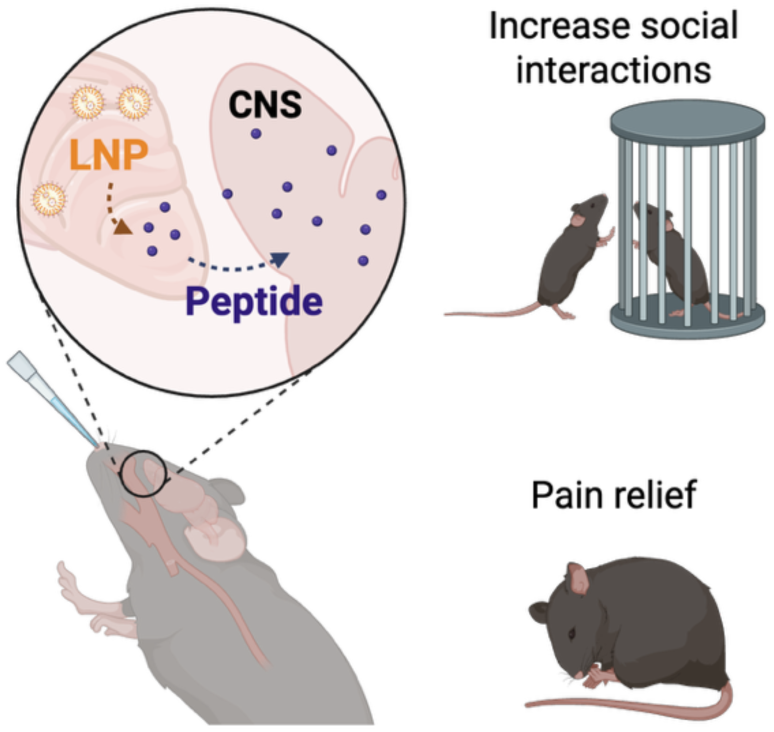

**One sentence summary:** Intranasal administration of mRNA-LNP enables local transfection in nasal epithelium and subsequent secretion of therapeutic oxytocin peptides into the brain, promoting social interactions and reducing pain.

## INTRODUCTION

The rapid development of effective COVID-19 (*1*) and cancer (*2*) vaccines has established mRNA-LNP therapeutics as new potent therapeutic class. While mRNA-LNP therapies targeting systemic or liver-based diseases have shown clinical efficacy(*3*), delivery of mRNA-LNP to the brain has been challenging (*4, 5*). Recent advances in brain targeting via intranasal delivery of therapeutics (*6*) or commensal bacterial expressing the neuroactive cytokine leptin (*7*) highlight the potential utility of delivering molecular-based therapeutics to the brain.

Pain is one of the most common neurological diseases, with chronic pain affecting 20-30% of the population (*8*). For neuropathic pain, which impacts 9% of the population, ∼70% of patients (hundreds of millions globally) do not respond to any available therapy and new therapeutics are desperately needed (*9, 10*). To address this critical need for non-opioid pain medicines, we developed a strategy to deliver mRNA-LNP encoded therapeutic cargo via an intranasal route using Generally Recognized as Safe (GRAS) status components (*11*), allowing for rapid therapeutic application.

## RESULTS

### Intranasal mRNA-LNP Uptake and Expression

Using the BioNTech COVID-19 mRNA backbone (*12*), we replaced the SARS-CoV-2 spike protein with luciferase (**Table S1**). The *in vitro* transcribed luciferase mRNA was then encapsulated in lipid nanoparticles (LNPs) comprised of various ionizable lipids plus DSPC/Cholesterol/PEG. 11 candidate LNP formulations were labelled with the lipophilic dye DiR and loaded with luciferase mRNA cargo, then administered intranasally. Following addition of luciferin, mice were imaged using IVIS (**Fig S1a**). While DiR signals were detected in all treated animals indicating effective LNP delivery (**Fig S1b,** quantified in **Fig S1c**), LNPs 3, 4, 5, 10, and ALC-0315 also drove detectable expression of the luciferase cargo (**Fig S1d,** quantified in **Fig S1e**).

Since ALC-0315 is a component of approved COVID-19 vaccines and has a well-known safety profile, we moved forward with ALC-0315-based LNP. Delivery of ALC-0315 LNP labelled with DiR and carrying luciferase mRNA was evaluated 24 hours post-inoculation. Again, DiR signal was detectable in the respiratory and olfactory epithelium of the nasal passage, and this led to localised and specific luciferase in the nasal passage with no detectable luciferase translation in the brain, liver, heart, lungs, spleen or kidney (**Fig S1f-g**). To further map mRNA-LNP target cells within the upper airways we delivered mRNA-LNP encoding *cre* to mice harboring a cre-inducible *tdTomato* reporter cassette (**Fig S2a**). 3 days after treatment, *cre* mRNA-LNP treated mice showed strong tdTomato signal in the respiratory (RE) and olfactory epithelium (OE) (**Fig S2b**). These signals were primarily in TUBA1A^+^ epithelial cells of the nasal epithelium (**Fig S2c**) and in epithelial cells overlapping or contiguous with the initial layers of OMP⁺ olfactory epithelium (**Fig S2d)**. No uptake was detected in CD45^+^ immune cells, CD68^+^ macrophages or Ly6G^+^ neutrophils (**Fig S2e-g**).

Having developed an LNP-mRNA system for effective intranasal delivery, we next sought to identify secreted factors that may have value as mRNA-LNP based pain therapeutics. For this, we leveraged high-resolution cell profiling data from animals in untreated and treated pain (*13*). Using single nuclei data from mice with entrenched neuropathic pain (spared nerve injury; SNI) or SNI mice treated for 21 days with iPSC-derived GABAergic painkiller neurons (iGABA) (*14*), we performed differential expression analysis to identify pathways that change after sustained analgesia. From these data we annotated upstream secreted peptides that can modulate these pathways, which include Orexin, GNRH, Netrin, EGF family and Oxytocin (**Fig 1a**). To evaluate an mRNA-LNP pain killer strategy we chose Oxytocin, a small cyclic peptide that has well-established roles in regulating social bonding (*15*) and reproduction (*16, 17*). Importantly delivery of intranasal oxytocin peptide can provide pain relief in both preclinical (*18–20*) and clinical settings (*21*). The clinical efficacy of oxytocin peptide has been inconsistent, due in part to the low efficiency of acute intranasal peptide delivery (*22*) and oxytocin’s short half-life (*23, 24*). We reasoned that the anatomical specificity and persistence of an intranasal mRNA-LNP delivery strategy may function as a long-lasting time-release oxytocin formulation, producing mRNA-encoded therapeutic peptide in the nose, which could then enter the brain where it can exert its neuroactive effects.

**Figure 1.**
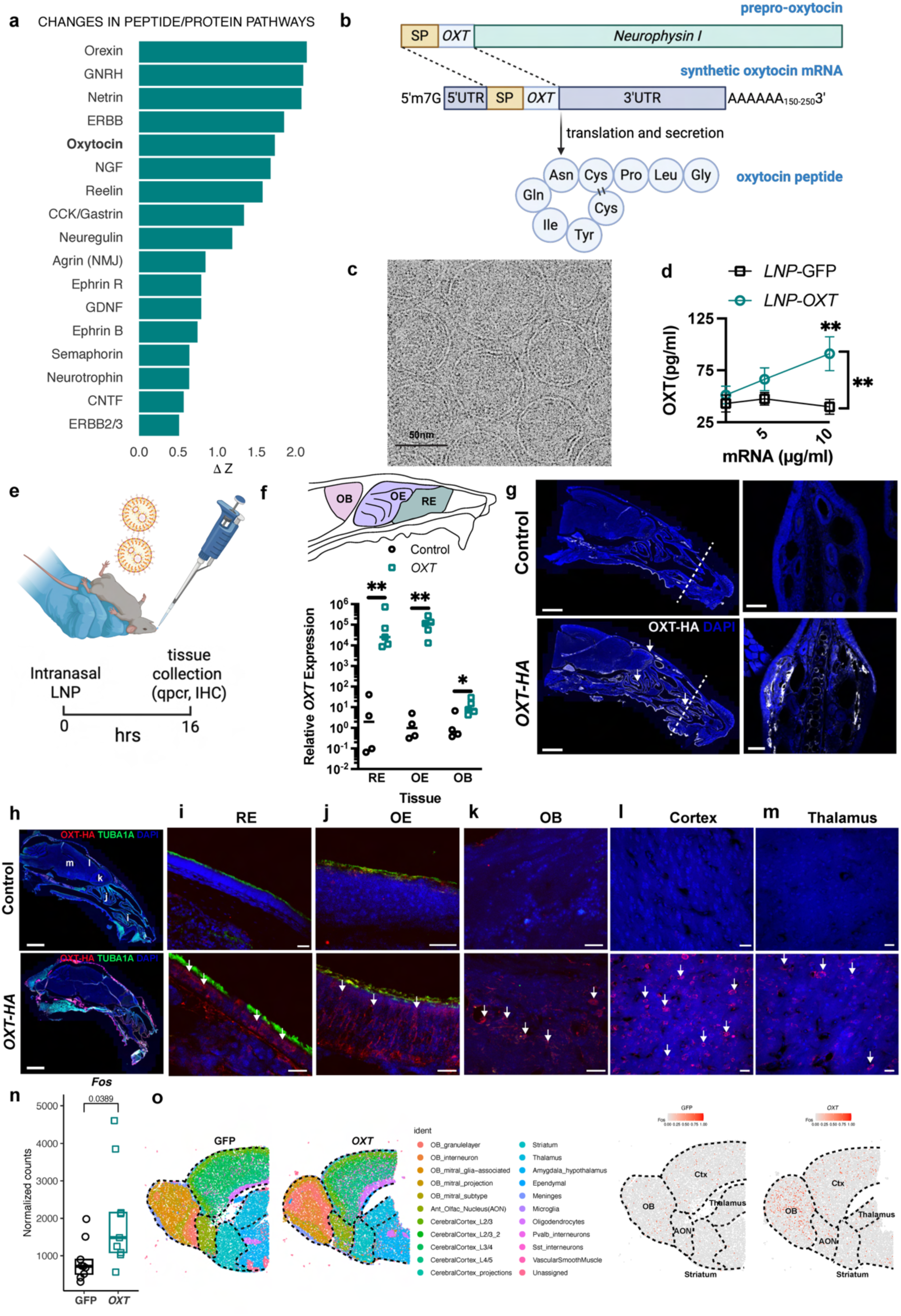
Assessing Oxytocin mRNA-LNP Uptake and Expression. **a**. Pathway analysis comparing SNI animals to iGABAergic transplant mice revealed changes in many nervous system pathways with secreted peptide pathways. **b.** An illustration depicting the synthesis strategy and translation of an mRNA strand into the oxytocin polypeptide. **c**. A CryoEM image of the lipid nanoparticles. **d**. Oxytocin peptide concentration in media collected 24 hours after adding OXT-LNP to HEPG2 cells. Data are presented as mean ± s.e.m. N = 3 biological replicates per condition. Significance was determined by two-way ANOVA,** p<0.01. **e.** Workflow for intranasal administration followed by tissue collection from the respiratory epithelium (RE). olfactory epithelium (OE). and olfactory bulb (OB). **f.** qPCR detection of exogenous human *OXT* mRNA in RE, OE and OB. Data are presented as mean ± s.e.m. N = 4 and 5 mice for control and *OXT* respectively. Significance determined by one-way ANOVA. p<0.05. p<0.01. **g**. Sagittal image (left panels, scale bar = 2 mm) of the control and OXT-HA skull with dashed lines indicating coronal sections (right panels, scale bar = 50 µm) where OXT-HA was detected in the RE. **h**. Immunohistochemistry images showing OXT-HA detection in the RE (**i**), OE (**j**), OB (**k**), cerebral cortex (**l) and thalamus (m).** Scale bars in h = 2 mm, i-m = 20 µm**. n.** *Fos* mRNA counts from anterior brain regions are higher in OXT treated mice, as determined via RNAseq. Data presented as mean ± s.e.m. N = 9, 3 per region (OB, Cortex, AON) per treatment. Statistical significance determined via unpaired t-test on normalised *Fos* counts of combined regions for each treatment. **o.** Spatial transcriptomics with 10X Genomics visium HD for control GFP and *OXT* treated tissue, confirming increased in *Fos* expression in anterior brain regions.

### Oxytocin Localization and Circuit Engagement

Native oxytocin transcript encodes an oxytocin / neurophysin I preprohormone that is processed to mature oxytocin in the magnocellular neurons of the paraventricular (PVN) and supraoptic (SON) nuclei of the hypothalamus (*17*). To bypass processing, we generated a synthetic oxytocin where the neurophysin I sequence is removed and the endogenous oxytocin signal peptide is fused directly to the mature oxytocin sequence, flanked by 5 and 3’ UTR/Poly A tail sequences used in the BioNTech COVID-19 vaccine, creating a 543 bp long mRNA strand (**Fig 1b, Table S1**). *In vitro* transcription of this construct lead to functional mRNA that could drive secretion of mature oxytocin in a dose-dependent fashion (**Fig S3a**), with maximum peptide secretion observed at 24 hours *in vitro* (**Fig S3b**). Notably, the human *OXT* mRNA sequence drove stronger OXT peptide production compared with the mouse sequence and was used for subsequent studies (**Fig S3a,b**). *Gfp* control or *OXT* mRNA was encapsulated in LNP and by zetasizer, *OXT* mRNA-LNP are ∼80 nm in diameter (**Fig S3c**) with zeta potential within an appropriate range (−5 mV; **Fig S3d**) for stable nanoparticles and cryo-electron microscopy confirmed uniform LNP morphology (**Fig 1c**). Importantly, *OXT* mRNA-LNP are functional and cells that receive *OXT* mRNA-LNP also produce OXT protein in a dose-dependent manner (**Fig 1d**).

Intranasal delivery of *OXT* mRNA-LNP (**Fig 1e**) led to a substantial increase in exogenous *OXT* mRNA levels in the respiratory epithelium (RE) and olfactory epithelium (OE), and modest increase in the olfactory bulb (OB) as measured by qPCR (**Fig 1f**). To detect exogenous OXT protein *in situ*, we added an hemagglutin (HA) tag to the OXT C-terminus (**Table S1**) and confirmed that OXT-HA is secreted after transfection (**Fig S3e**). 24 hours after intranasal *OXT-HA* mRNA-LNP administration, we assessed OXT-HA protein biodistribution by immunofluorescent microscopy. Compared to control, we observed strong and localised OXT-HA protein in the mouse head across the respiratory epithelium, olfactory epithelium, and in the brain (**Fig 1g**, see arrows). Co-staining with the epithelial marker TUBA1A (*25*) showed OXT-HA co-localization in both the respiratory and olfactory epithelium **(Fig 1h-j**, see arrows). OXT-HA was also observed in multiple regions of the brain including the olfactory bulb, cortex, and thalamus **(Fig 1k-m**, see arrows). To assess OXT peptide functional engagement in the CNS we quantified *Fos* expression, a rapid neural activation marker upregulated by oxytocin(*26, 27*). RNA-seq showed that intranasal delivery of *OXT* mRNA-LNP (compared to *Gfp*-mRNA-LNP control) drove significant *Fos* upregulation in anterior brain regions (**Fig 1n**) and spatial transcriptomics highlighted robust *OXT* mRNA-LNP-driven *Fos* activation in forebrain regions implicated in social behaviour, pain and sensory integration (**Fig 1o**, **Fig S4a-e**). Notably, thalamic nuclei (which control relay of sensory information including pain) showed the strongest *Fos* increase (**Fig 1o**, **Fig S4f**). Cortical layers (L2–L5) and striatal regions, linked to pain processing, were also engaged, alongside parvalbumin and somatostatin interneurons, which set inhibitory tone within pain-processing networks (**Fig S4f)**. *Fos* activation was also observed in the olfactory bulb, the anterior olfactory nucleus, the amygdala, and the hypothalamus, brain regions that are central to social behaviour downstream of OXT.

### Social and Safety Effects of Intranasal mRNA-LNP Therapy

We next tested if intranasal *OXT* mRNA-LNP could alter animal behaviour. Since OXT is best known to promote social interactions(*17*), we first evaluated social behaviour using the three-chamber test. Before *OXT* dosing, animals were acclimated to the test and sorted for comparable baseline social behaviour between groups (**Fig 2a**). Animals were then rested for 2 days, treated with an escalating dose (0, 20, 40 and 80 µg/kg) of intranasal *OXT* mRNA-LNP and then tested for social behaviour. Animals displayed a U-shaped dose response in increasing social interactions, with peak interactions at 40 µg/kg (**Fig 2b**). To rule out LNPs as a contributor to the increased social interactions, we assessed the effects of *Gfp* mRNA-LNP vs *OXT* mRNA-LNP. Animals dosed with *Gfp* mRNA-LNP showed no preference for social interaction, as expected with repeated exposures to this test. By comparison, animals that received *OXT* mRNA-LNP displayed significantly enhanced social behaviour (**Fig 2c**) and this effect was observed in both males (**Fig 2d**) and females (**Fig 2e**). *OXT* mRNA-LNP did not impact overall motor function as assessed by rotarod (**Fig 2f**). Additionally, using metabolic chambers we did not observe any impact of *OXT* mRNA-LNP dosing on general activity, food intake, body weight or respiration (**Fig 2g**). Finally, ALC-0315 based LNP can promote inflammation in some contexts (*28*), however histological analysis of the respiratory and olfactory epithelia showed no significant increase in inflammatory cell recruitment or tissue damage after treatment (**Fig 2h-j**). Overall, these data show that intranasal mRNA-LNP treatment is a safe and effective method to achieve sustained (>24 hour) and physiologically relevant delivery of bioactive peptides like OXT to the brain.

**Figure 2.**
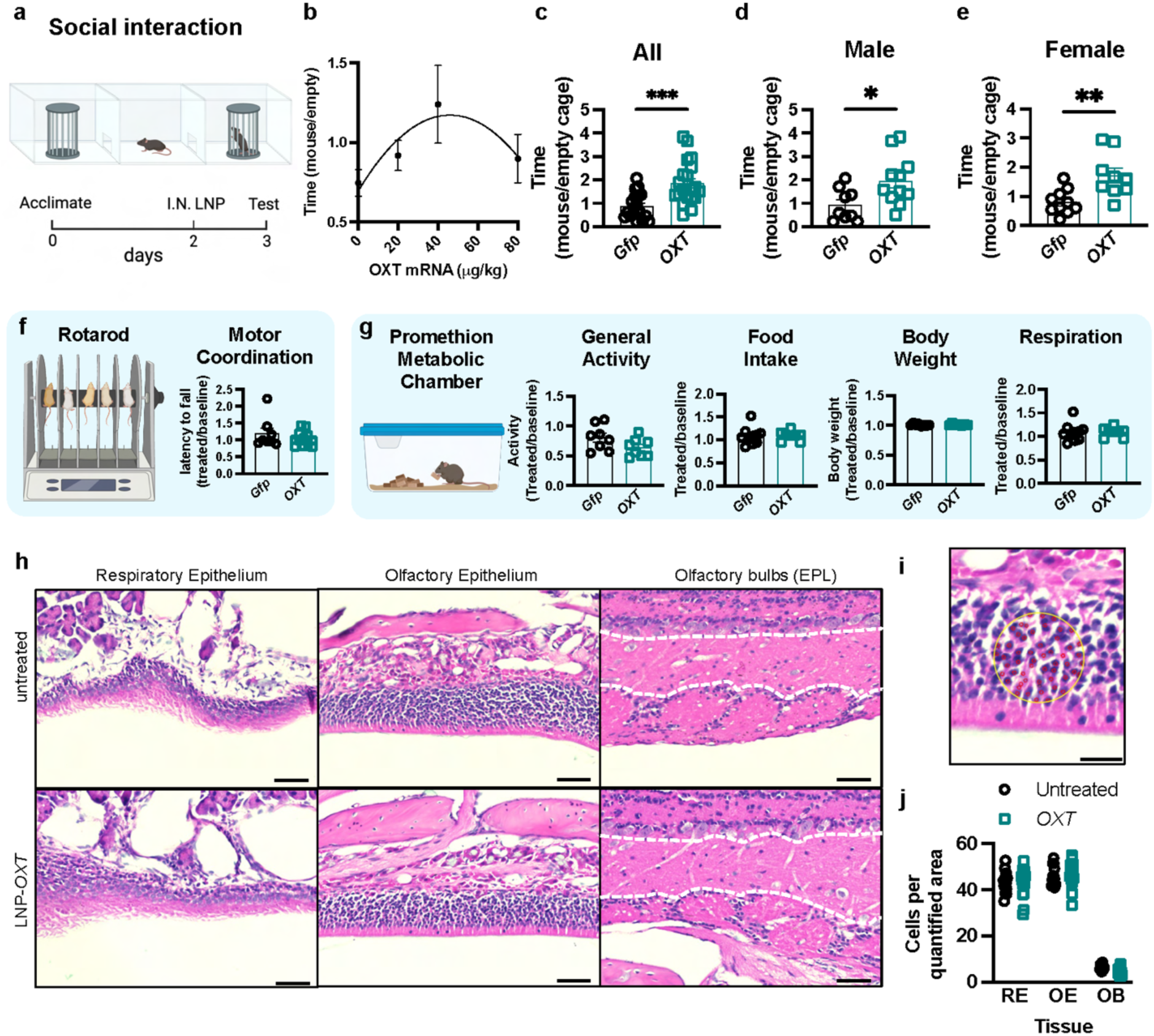
OXT-LNP increase social interaction without inducing side effects or inflammation. **a.** Illustration of social interaction experimental setup. **b.** OXT-LNP increases social interaction in a dose dependent manner, peaking at 40 µg/kg. Data presented as mean ± s.e.m. N = 6-8 per dose. **c.** Further experiments in male (**d**) and female (**e**) mice with 40 µg/kg mRNA showing increased time spent with a mouse enclosure after OXT-LNP administration. **f.** Rotarod test showing no motor function impairment. **g**. Metabolic cage data showing no changes in general activity. food intake. body weight. or respiration. **h.** H&E histology images of olfactory bulb (EPL), olfactory epithelium, and respiratory epithelium from untreated and OXT-LNP-treated mice. Scale bars = 50 µm **i**. A representative image of a quantified area. Scale bars = 20 µm. **j**. Quantification of cell nuclei per area. showing no significant increase in cell numbers. All data are presented as mean ± s.e.m. Significance was determined by t-test. with *p<0.05, **\*\***p<0.01, **\*\*\***p<0.005.

### Analgesic Effects Across Pain Models

To evaluate the utility of *OXT* mRNA-LNP as an analgesic, we dosed animals with LNP and then assessed their behaviour in a series of pain paradigms. In healthy uninjured mice, *OXT* mRNA-LNP significantly reduced baseline mechanical sensitivity (**Fig 3a,b**) without affecting thermal responses (**Fig 3c,d**). Furthermore, in a post-operative pain model (**Fig 3e**), *OXT* mRNA-LNP treatment significantly reduced mechanical pain **(Fig 3f**), with an efficacy comparable to the anti-inflammatory drug Meloxicam **(Fig 3g**). In neuropathic (SNI) mice (**Fig 3h**), we found intranasal *OXT* mRNA-LNP provided relief from mechanical **(Fig 3i**), heat **(Fig 3j**) and cold hypersensitivity (**Fig 3k).** We confirmed the analgesic activity of *OXT-*mRNA-LNP using the “Blackbox,” an objective behavioural system equipped with a near-infrared (NIR) vision camera and machine learning analysis to quantify luminance ratios between injured and uninjured paws. This ratio reflects the relative surface contact area of each paw, serving as a proxy for weight-bearing and mechanical pain intensity. SNI mice typically reduce contact on the injured paw (**Fig 3l**), resulting in a luminance ratio of ∼0.2 (i.e., the injured paw accounts for only 20% of total contact; **Fig 3m**). Importantly, treatment with *OXT* mRNA-LNP provided significant analgesia (**Fig 3n**), and this effect was similar to the standard-of-care Gabapentin (**Fig 3o**), however the effects of *OXT-*mRNA-LNP last >24 hours per dose while the effects of Gabapentin wear off after ∼4 hours. Of note, while *OXT* mRNA-LNP promoted social behaviour in both male and female mice, the analgesic effects were only observed in male mice (**Fig S5**). Overall, we report the surprising finding that intranasal mRNA-LNP can be used to deliver secreted neuroactive peptides to the brain, unlocking a new mRNA-based therapeutic platform to treat pain and other major neurological diseases.

**Figure 3.**
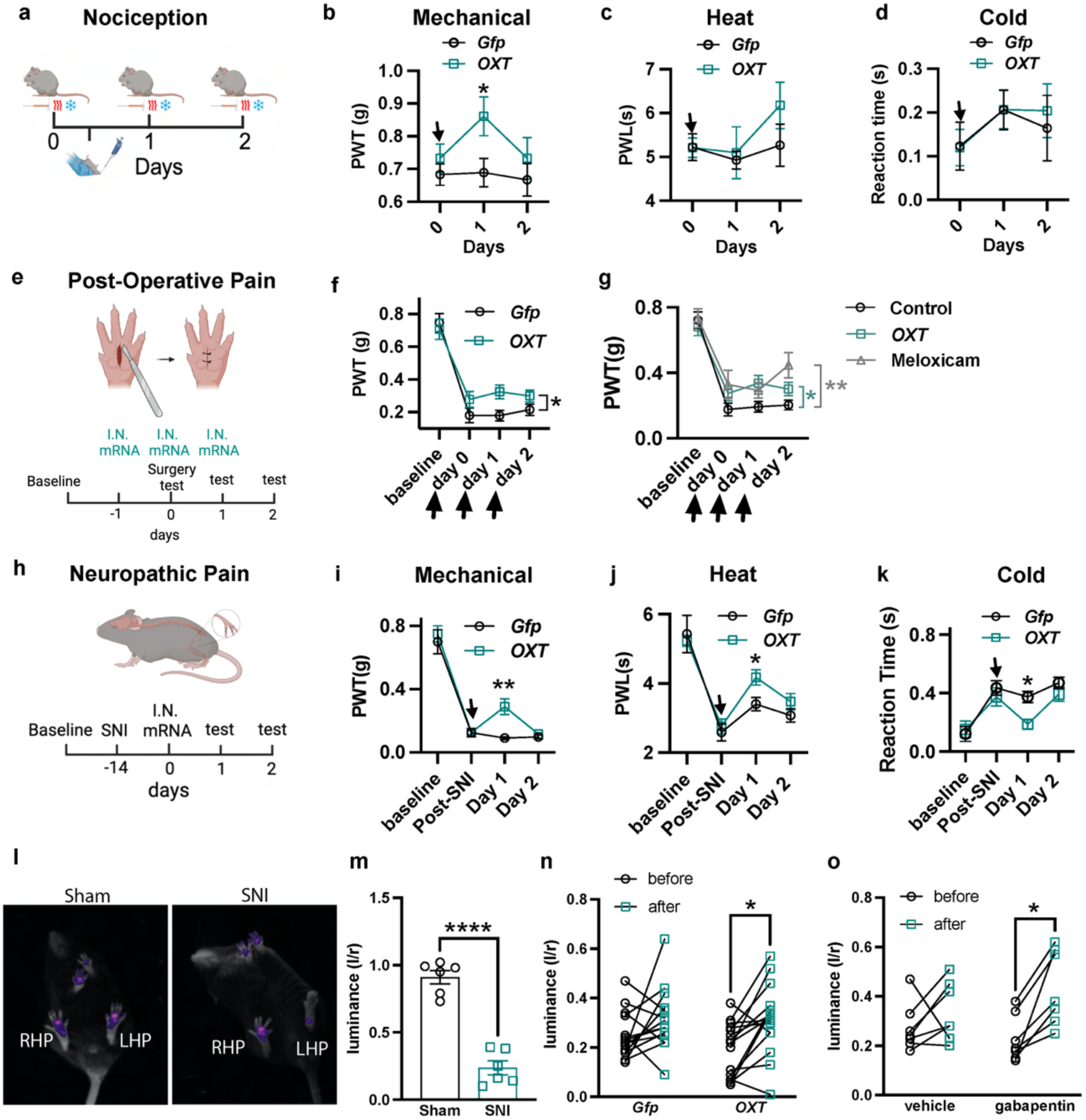
Analgesic Effects of OXT-LNP in Pain Models. **a.** Illustration of the nociception study design. Mechanical (**b**), heat(**c**) and cold (**d**) sensitivity in healthy mice after administration. Data presented as mean ± s.e.m, n = 12 per treatment. Significance was determined by two-way ANOVA vs. GFP control, *p<0.05. **e.** Illustration of the post-operative pain model. **f.** Mechanical pain in mice treated with *<u>OXT</u>* or GFP controls. **g.** Comparison of *OXT* pain relief with Meloxicam. Data are presented as mean ± s.e.m, n = 10 per treatment. Significance was determined by two-way ANOVA vs. GFP control,*p<0.05,**p<0.01 or one-way ANOVA vs. vehicle control. **h.** Illustration of the neuropathic pain study. Mechanical (**i**), heat (**j**) and cold (**k**) sensitivity in neuropathic mice after administration. Data are presented as mean ± s.e.m., n = 8 per treatment. Significance was determined by two-way ANOVA vs. GFP control, *p<0.05, **p<0.01. **l.** Representative luminance images of naturalistic behaviour using the automated quantitative imaging with Blackbox. **m.** SNI mice display decrease in left hind paw pressure (L/R luminance). **n.** *OXT* produced significant analgesia in the injured paw, similar to the first-in-class treatment for neuropathic pain, Gabapentin (**o**). Data are presented as mean ± s.e.m. Significance was determined by paired multiple T-test comparisons (*p<0.05).

## DISCUSSION

Our findings demonstrate that intranasal delivery of mRNA-LNP encoding secreted neuroactive peptides represents a viable and non-invasive strategy to target the central nervous system (CNS) with peptide therapeutics. Historically, the blood–brain barrier (BBB) has posed a major obstacle for peptide-based therapeutics, limiting their clinical utility. Intranasal administration of mRNA-LNPs results in mRNA uptake and expression in the nose, where oxytocin peptide is then produced and enters the brain. This approach aligns with prior evidence supporting nasal delivery of small peptides but extends the concept by introducing a genetic blueprint for sustained local production rather than transient peptide diffusion. Our findings complement recent work using engineered commensals for nose-to-brain delivery (*7*), highlighting the growing interest in leveraging nasal pathways for CNS-targeted biologics.

Oxytocin protein localized to brain regions including the olfactory bulb, anterior olfactory nucleus, cortex and thalamus, areas implicated in sensory integration, social processing, and reward. Our data aligns with radioligand mapping of oxytocin receptor sites in the forebrain, supporting the functional engagement of circuits implicated in social and sensory processing (*22*). This suggests functional engagement of circuits beyond nociception, consistent with oxytocin’s established role in social bonding and affective regulation. Behavioural assays support this, revealing a U-shaped dose-response in social interaction, a phenomenon widely reported in oxytocin literature and likely reflecting receptor saturation or feedback mechanisms (*29*). This U-shaped profile may serve as a self-limiting mechanism, providing an additional layer of control that could reduce reliance or addictive potential, an important consideration for therapeutics targeting reward pathways.

The *Fos* signature points to a network-level analgesic effect, engaging thalamocortical and limbic circuits while recruiting cortical inhibitory interneurons, a systems architecture distinct from peripheral nociceptor silencing or spinal-only analgesia. Intranasal *OXT* mRNA-LNP produced consistent analgesia across multiple modalities and pain paradigms. The magnitude of pain relief observed was comparable to standard-of-care agents such as meloxicam and gabapentin yet long lasting and delivered via a non-invasive route without systemic side effects or inflammation. This dual action of enhancing social behaviour and reducing pain may be particularly valuable for patients with chronic pain, who often experience social isolation that impacts pain disease (*30*). Notably, in mice the pro-social effects of *OXT* mRNA-LNP were observed in both sexes, however the analgesic effects were only observed in male mice. This differs from intranasal OXT effects in humans, where analgesia was observed in both male and female recipients (*31*) and the gender-specific OXT activity on pain we observe may be a rodent-specific phenomenon.

Collectively, these results establish intranasal mRNA-LNP delivery as a promising platform for CNS-targeted therapeutics. By combining anatomical accessibility with genetic programmability, this approach offers a scalable and patient-friendly alternative to invasive procedures or short-lived peptide sprays. Future work should explore long-term clinical safety, and applicability to other neuroactive peptides or proteins. If successfully translated, this strategy could redefine treatment paradigms for pain and other major neurological disorders.

## MATERIALS AND METHODS

### Cloning of mRNA Template and *In vitro* Transcription

Firefly Luciferase, Human and mouse oxytocin mRNA vectors were constructed by sequentially assembling the corresponding gene gBlocks (Integrated DNA Technologies) into a Pfizer mRNA backbone plasmid using PCR products and the NEBuilder HiFi DNA Assembly Kit (New England Biolabs). The assembled plasmids were verified by Sanger sequencing, followed by linearization with the BspQI restriction enzyme (NEB, #R0712L). Sequences are provided in Table S1. mRNA synthesis was then performed using the HiScribe® T7 High Yield RNA Synthesis Kit (New England Biolabs) and N1-Methylpseudouridine-5’-Triphosphate (TriLink BioTechnologies, N-1081).

### *In vitro* mRNA Assessment

HEK 293T cells were seeded at a density of 3 × 10⁵ cells per well in a 12-well plate one day prior to transfection. Transfection of human and mouse mRNA was carried out using Lipofectamine™ MessengerMAX™ reagent (Thermo Fisher Scientific). Culture media were collected at 24, 48, and 72 hours post-transfection, and oxytocin concentrations were measured using the Oxytocin ELISA kit (ENZO Life Sciences). For LNP experiments, HEPG2 cells were seeded in 12-well plates at 0.3 × 10⁶ cells per well and cultured for 24 h. ApoE solution (20 µL/mL) was mixed with LNP–mRNA suspension at a 1:1 volume ratio and incubated at room temperature for 15–30 min to allow ApoE association. The mixture was then diluted with DMEM to a final volume of 500 µL and added to cells after aspirating the culture medium. Cells were incubated for 24 h before collection for protein extraction.

### ELISA

Conditioned media collected at 24-, 48-, and 72-hours post-transfection were processed for oxytocin quantification. Proteins were extracted using HyperSep™ C18 Cartridges (3 ml, Thermo Fisher Scientific), and the eluates were evaporated using a Genevac EZ-2 Centrifugal Evaporator (SP Scientific, Ipswich, UK). The dried samples were reconstituted in 110 µl of the assay buffer provided with the ELISA kit, and 100 µl of the reconstituted solution was used for measurement. Oxytocin concentrations were quantified using the Oxytocin ELISA Kit (ADI-900-153, Enzo Life Sciences) according to the manufacturer’s instructions. The assay has a detection limit of ∼15 pg/ml and an upper quantification limit of 1000 pg/ml.

### Lipid Nanoparticle (LNP) Encapsulation and Quality Control

Firefly *Luc* mRNA was encapsulated into lipid nanoparticles (LNPs) with DiR for in vivo administration and biodistribution tracking. 11 ionizable lipids were selected based on *in vitro* screening. These include ALC-0315 and several proprietary candidates designed to enhance absolute protein expression and minimize cytotoxicity using endogenous and biodegradable motifs. Full structural details will be disclosed following patent publication; all functional data supporting their performance are provided herein. The mRNA and lipid components – ionizable cationic lipid (50.1%), DSPC (distearoylphosphatidylcholine, 10.6%), cholesterol (37.2%), ALC-0159 (PEG-lipid, 1.9%), and DiR (fluorescent dye, Thermofisher, D12731, 0.2%) were mixed using a microfluidic mixer (NanoAssemblr™ Ignite™ nanoparticle formulation system, Cytiva) at a target molar charge ratio of 6:1 (cationic lipid : mRNA). Following encapsulation, the LNP solution was buffer exchanged with 10x volume of 0.01 M Tris buffer and processed using an Amicon® Ultra-15 Centrifugal Filter. The mixture was centrifuged at 2000 × g to remove ethanol from the solution. The remaining mRNA-LNP complexes are stored in 0.01 M Tris with 10% sucrose. The resulting mRNA-LNP complexes were subjected to quality control assessments. Encapsulation efficiency was determined using the Quant-iT™ RiboGreen RNA Assay Kit (Invitrogen™). The size distribution and zeta potential of the LNPs were analyzed using a Zetasizer instrument (Malvern analytical).

### IVIS Imaging

For LNP biodistribution studies, mice were administered intranasally with DiR-labeled mRNA-LNP formulations under light anesthesia. Once mice exhibited no reflexive responses (respiratory rate ∼144 breaths/min), they were gently scruffed, positioned supine, and the formulation was pipetted into both nostrils. At 16 hours post-administration, tissue was collected from animals and incubated in 500 µM D-luciferin in 1X PBS and imaged using an IVIS Spectrum system (PerkinElmer). Fluorescence (DiR) and Luminescence (Luciferin) Images were acquired using auto-exposure settings with binning factor 8 and analyzed using Living Image software (v4.7.3). Regions of interest (ROIs) were drawn over the nasal cavity and quantified as radiant efficiency (photons/sec/cm²/sr per µW/cm²). Sample sizes for each condition are indicated in figure legends.

### Social Assessment

The three chamber social interaction test was performed as previously described(*32*). The apparatus consisted of a white enclosure divided into three chambers: two end chambers, each containing a wire cage (10 cm diameter) and a central chamber where the animal was initially placed at the start of each test. Mice could move freely between chambers through openings in the dividers separating chambers.

Mice were firstly habituated to the empty apparatus for 10 min. 1 hr later, the sociability test was performed. One wire cage remained empty while the other contained an age, sex and genetic-background matched stimulus animal. The test mouse was placed in the central chamber and allowed to freely explore for 10 min. All movement was recorded from above using a Webcam. Locomotor activity and the time spent investigating each cage was analysed using Any-MAZE tracking software (Version 7.20)

The social interaction test was performed twice. The first session served as a baseline to assign animals to *Gfp* or *OXT* treatment groups, ensuring comparable averages and variation between groups. For initial dose-response studies, female mice were with 0, 20, 40 and 80 µg/kg of *OXT* while comparisons between *Gfp* and *OXT* groups in males and females were done at 40 µg/kg. The second session was performed 24 hr following intranasal administration of mRNA-LNP.

### *In vivo* Nociceptive Assessment

To evaluate the effects OXT-LNP, the formulation was administered intranasally at 40 µg/kg to C57BL/6 male mice (9-18 weeks old). Following habituation to the behavioral testing apparatus and environment, the animals underwent pain behavior assessments, including the von Frey test, Hargreaves test, and acetone test. These tests were conducted at three time points: before administration, and at 24- and 48-hours post-administration.

### Post-operative Pain Model

Post-surgical pain was induced using a plantar incision model as previously described (*33*). Briefly, mice were anesthetized with isoflurane (2–3%) and a 2 mm longitudinal incision was made through the skin and fascia of the plantar surface of the left hind paw. The underlying muscle was elevated without transection. The wound was closed with 5-0 nylon sutures and animals were allowed to recover in warmed cages. 3 intranasal administrations of 40 µg/kg *OXT* mRNA-LNP or control *Gfp* mRNA-LNP were performed on the day before surgery, 24- and 48-hours post-surgery under light anesthesia. Mechanical sensitivity was assessed using von Frey filaments at baseline and at 24, 48, 72 and 96 hours after dosing.

### Spared Nerve Injury (SNI) Model

Neuropathic pain was induced using the spared nerve injury model(*34*). Under isoflurane anesthesia, the sciatic nerve branches were exposed, and the tibial and common peroneal nerves were tightly ligated and transected, leaving the sural nerve intact. The muscle and skin were closed in layers. Animals were monitored daily for recovery. Behavioral testing (von Frey, Hargreaves, acetone) was performed at baseline and 14 days post-surgery to confirm neuropathic hypersensitivity. Intranasal dosing of 40 µg/kg *OXT* mRNA-LNP or *Gfp* controls was performed under light anesthesia, and pain assessments were repeated at 24 and 48 h post-treatment.

### Tissue Preparation for H&E and Immunostaining

Mouse heads and brains were immersion-fixed in 4% paraformaldehyde (PFA) overnight at 4 °C, followed by decalcification in UltraPure™ 0.5 M EDTA, pH 8.0 (Invitrogen) for 5 days at 4 °C with daily solution changes. Samples were dehydrated in 30% sucrose overnight at 4 °C, embedded in Tissue-Tek® O.C.T. compound (Sakura Finetek), and cryosectioned at 30 µm thickness using a cryostat.

### Immunostaining

Sections were blocked in PBS containing 5% normal goat serum (NGS) and 0.3% Triton X-100 for 30 min at room temperature. Primary antibodies were diluted in the same blocking solution and applied overnight at 4 °C at the following dilutions: 1:200 for α-Tubulin (T6199, Sigma), 1:200 for CD45 (103111, BioLegend), 1:200 for CD68 (14-0681-82, eBioscience), 1:200 for HA tag (ab9110, Abcam), 1:300 for Olfactory Marker Protein (ab183947, Abcam), 1:200 for Ly6G (RB6-8C5, eBioscience), 1:200 for EpCAM (118214, BioLegend), and 1:200 for GFP (A10262, Life Technologies). After washing, fluorophore-conjugated secondary antibodies (Thermo Fisher Scientific) were applied for 2 h at room temperature in the dark as follows: goat anti-mouse Alexa Fluor 488 (A-11029, 1:1000), goat anti-mouse Alexa Fluor 647 (A-21235,1:1000), goat anti-rabbit Alexa Fluor 488 (A-11008, 1:1000), goat anti-rabbit Alexa Fluor 555 (A-21428, 1:1000), and goat anti-rabbit Alexa Fluor 647 (A-21245, 1:1000), goat anti-rat Alexa Fluor 647 (A-21247, 1:1000), goat anti-guinea pig Alexa Fluor 647 (A-21450, 1:1000).

Sections were mounted using ProLong™ Gold Antifade Mountant with DAPI (Thermo Fisher Scientific, P36931). Confocal images were acquired using a Nikon C2 confocal microscope, and whole-slide images were scanned with a Zeiss Axio Scan.Z1 slide scanner. Image analysis was performed using QuPath (v0.4.3).

### H&E Staining

Hematoxylin and eosin (H&E) staining was performed on paraffin-embedded tissue sections using standard procedures. Sections were deparaffinized, rehydrated through graded ethanol solutions, and brought to water before staining with Harris’s hematoxylin for 30 s. Slides were rinsed in water, differentiated in acid alcohol with 10 rapid dips to remove excess stain, and washed again in water. Sections were blued in Scott’s bluing solution for approximately 7–8 s, washed in water, and examined microscopically to confirm clear nuclear staining with minimal cytoplasmic staining. Slides were then washed in water for 3 min, briefly rinsed in 70% ethanol for 30 s, and counterstained twice in eosin. Following eosin staining, sections were dehydrated through graded alcohols, cleared in two changes of xylene for 2 min each, and mounted with a permanent mounting medium.

### RNAseq

Brain tissue from OB, Cortex and AON of *Gfp*- and *OXT*-LNP mice were collected 16 h post administration and RNA was extracted using Bioline ISOLATE II kit and quantified with nanodrop before sequencing at 30 million paired-end reads with Novogene AIT (Singapore) using the NovaseqXPlus. Sequenced data was aliqned using Salmon and analysed with DESeq2. Normalised counts were then extracted for Fos for statistical analysis.

### Tissue Preparation and Sectioning for CytAssist (Visium HD, 10X Genomics)

FFPE blocks were prepared from mouse brain tissue post-fixed in 4% paraformaldehyde (PFA) for 24 h. Sagittal sections (5 µm) were collected from the median plane and mounted on glass slides for Visium HD and CytAssist workflows. Visium CytAssist Spatial Gene Expression for FFPE was performed on a subset of samples following the *Demonstrated Protocol for Deparaffinization, H&E Staining, Imaging & Decrosslinking* (CG000520) and processed using the *Visium CytAssist Spatial Gene Expression Reagent Kits User Guide* (CG000495). Alignment and segmentation were performed using Space Ranger v4.0, and downstream analysis followed the same Seurat-based pipeline described above.

### Visium HD Spatial Gene Expression

H&E staining and imaging were performed according to the *Visium HD FFPE Tissue Preparation Handbook* (CG000684). Subsequent processing and sequencing followed the *Visium HD Spatial Gene Expression Reagent Kits User Guide* (CG000685). Alignment and feature calling were performed using **Space Ranger v3.0**, which outputs data at native 2-µm resolution and binned resolutions of 8 µm and 16 µm. Downstream analyses were conducted on 16-µm bins. This resolution balances computational efficiency with robust cell-type annotation at near single-cell scale. UMI count matrices from *Gfp* and *OXT* sections were normalised individually with before merging using Seurat V5, sampling 20% of the dataset (*35*). Variable features were identified, data scaled, and PCA performed (30 PCs). Graph-based clustering was applied (resolution = 0.8) to define cell types. Clusters were determined via differential gene expression (DGE) analysis and manual annotation using known marker genes (Table S2). Final annotations were propagated to the full dataset. ImageFeaturePlot was used to plot normalised Fos counts, and dotplot was used to show average expression profile between the treatments for each cluster.

### Blackbox Assessment and Analysis

Sham or SNI mice (left hindpaw injury) were habituated on the Blackbox platform for 30 min the day before baseline recording (15 min). Baseline left/right hindpaw luminance was averaged and analyzed using Blackbox software (v0.1.1) to assign treatment groups evenly. For mRNA dosing, mice received intranasal administration under light anaesthesia and were recorded 16 h later. Vehicle and gabapentin were administered intraperitoneally 30 min before recording. Post-treatment luminance was compared to baseline and analyzed using paired t-tests

### Promethion Metabolic Monitoring

Mice were habituated in Promethion metabolic chambers (Sabre System International) for 12 h prior to recording. Baseline weight, food, and water intake were measured to confirm consistency with system readouts. Animals were monitored for three light/dark cycles, then administered intranasally with 40 µg/kg *Gfp* or *OXT* mRNA-LNP. Following dosing, mice were returned to their chambers and recorded for an additional three light/dark cycles. Data on activity, food intake, body weight, and respiration were extracted using OneClickMacro (v2.53.2) and normalized to baseline.

### Rotarod

Motor coordination was assessed using an accelerating rotarod (4-40 rpm over 5 min). Mice underwent four trials without prior training, with animals from the same cage tested together (up to four per run). After positioning at 4 rpm, the rod was ramped to 40 rpm over 300 s, and latency to fall was recorded automatically or manually. Trial termination reasons (falling, jumping, passive rotation) were noted. Padding was placed beneath the rod to minimize injury risk. The procedure was repeated the day after intranasal administration, and latency to fall was normalized to baseline.

## Acknowledgments

We acknowledge the University of Sydney core facilities: Sydney Microscopy, Laboratory Animal Services, Mass Spectomery, Sydney Analytical, Sydney Imaging and Sydney Informatics Hub, for their assistance with CryoEM imaging, animal maintenance, peptide identification, LNP QC via zetasizer, IVIS imaging, and access to Ingenuity Pathway Analysis, respectively. We thank the Charles Perkins Centre and Dr John and Anne Chong for their continuous support of our work. We thank Cytiva/Precision Nanosystem for their guidance in LNP encapsulation.

## Author Contributions

Conceptualization: L.L and G.N. Ionisable lipid production: R.O. and C.F. mRNA and LNP production: S.C., R.C., K.F., L.L. Encapsulation: F.S., P.T., R.C., K.F., L.L. *In vitro* assays: L.L., K.F., R.C., M.B. Peptide Detection: L.L., K.F., M.B., R.C, L.J.M. *In vivo* assessments: L.L., K.F., M.B., T.D. Funding acquisition: L.L, G.N. Supervision: L.L. and G.N. Writing original draft: G.N. Review and editing: L.L., M.B., P.T., C.F., R.C. and G.N.

## Data, code, and materials availability

All data in the main text or the supplementary materials are available at dryad (DOI: 10.5061/dryad.gf1vhhn3k).

---

**Fig S1.**
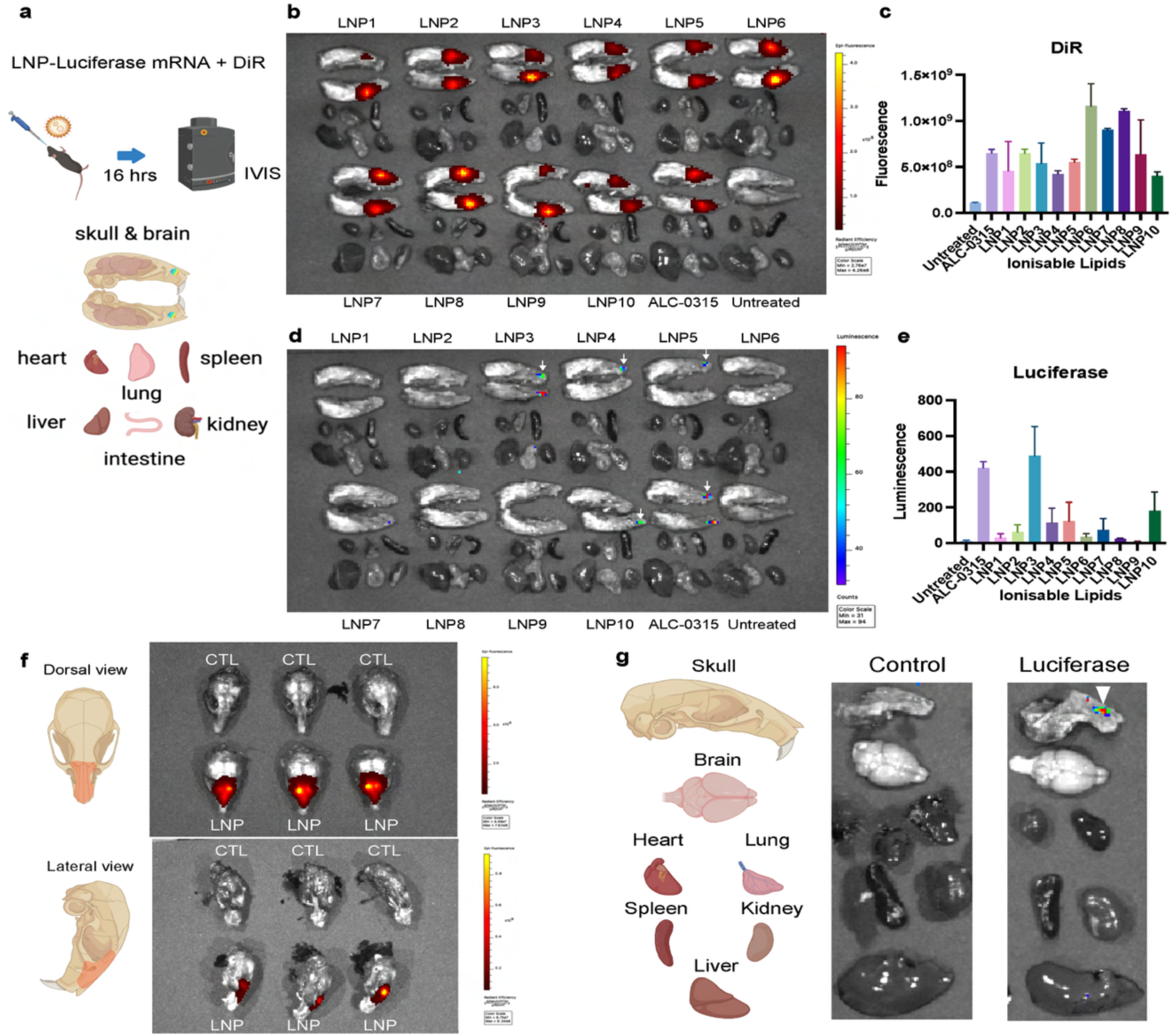
Intranasal mRNA-LNP Delivery to the CNS. **a** Intranasal administration of 40 ug/kg of Luciferase mRNA in DiR-containing LNPs followed by IVIS imaging. **b.** DiR distribution for all 11 candidate LNPs in the nasal passage (n=1 biological replicate per LNP, 2 nasal regions quantified per animal) **c.** Quantification of DiR signals. **d.** High luminescence in the nasal passage for LNP3, 4, 5, 10 and ALC-0315. **e.** quantification of luciferase luminescence. **f**. Confirmation of DIR uptake in nasal passage following intranasal ALC-0315 LNP (n=3). **g.** Luciferase expression confirmed in nasal passage following intranasal ALC-0315 administration.

**Fig S2.**
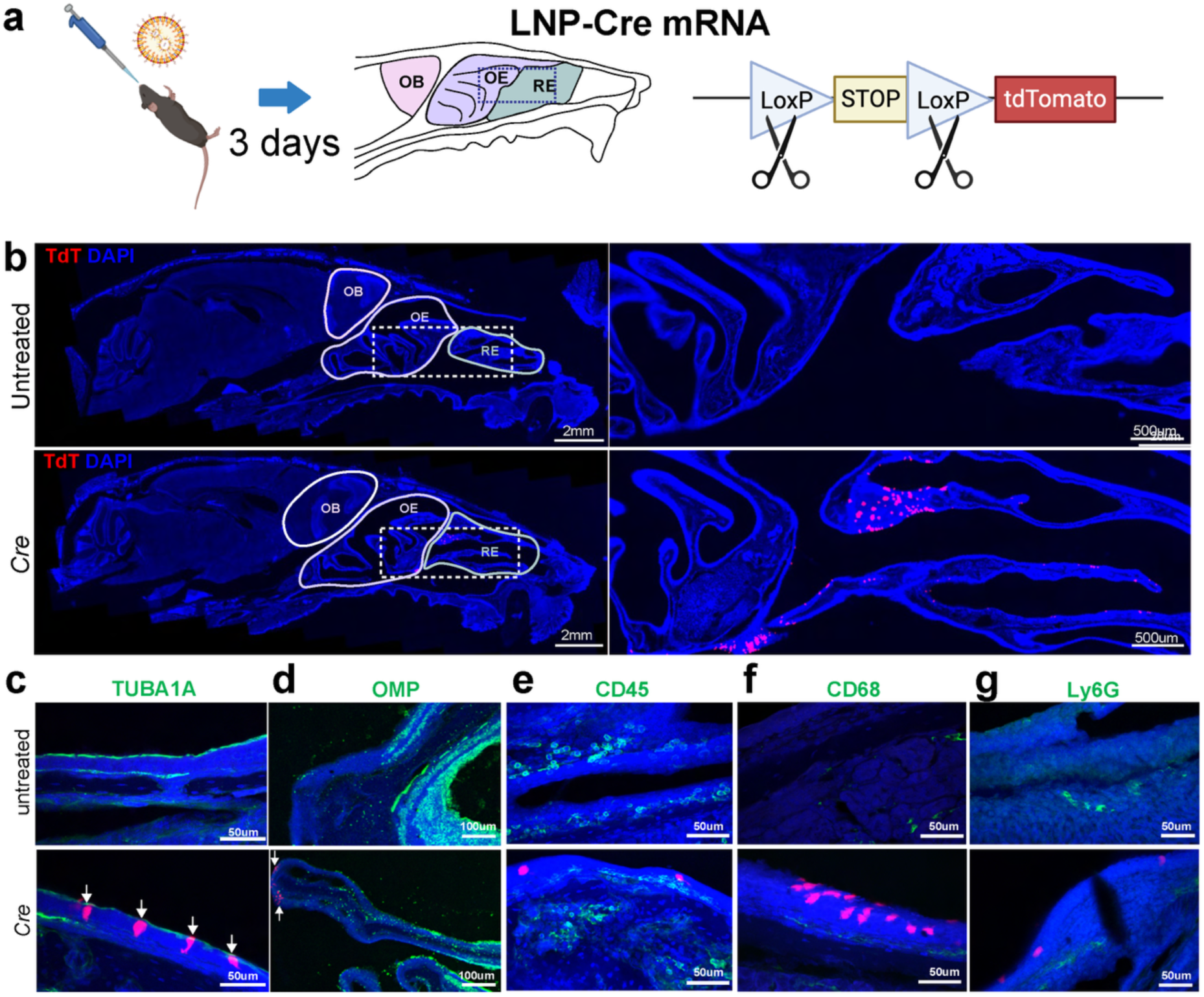
Epithelial cells take up mRNA-LNP. **a.** Workflow for Cre-mediated recombination in an Ai9-tdTomato reporter mouse. **b.** Tiled images of Control and *Cre*-treated Ai9 Tdtomato mice, showing expressing in RE and OE. **c**. TdTomato fluorescence in the TUBA1A^+^ respiratory (RE) and **d.** OMP^+^ olfactory epithelium (OE) mice that received LNP-Cre mRNA (Cre). No Tdtomato signals in CD45+ immune cells (**e**), CD68+ macrophages (**f**), or Ly6g+ neutrophils (**g**).

**Fig S3.**
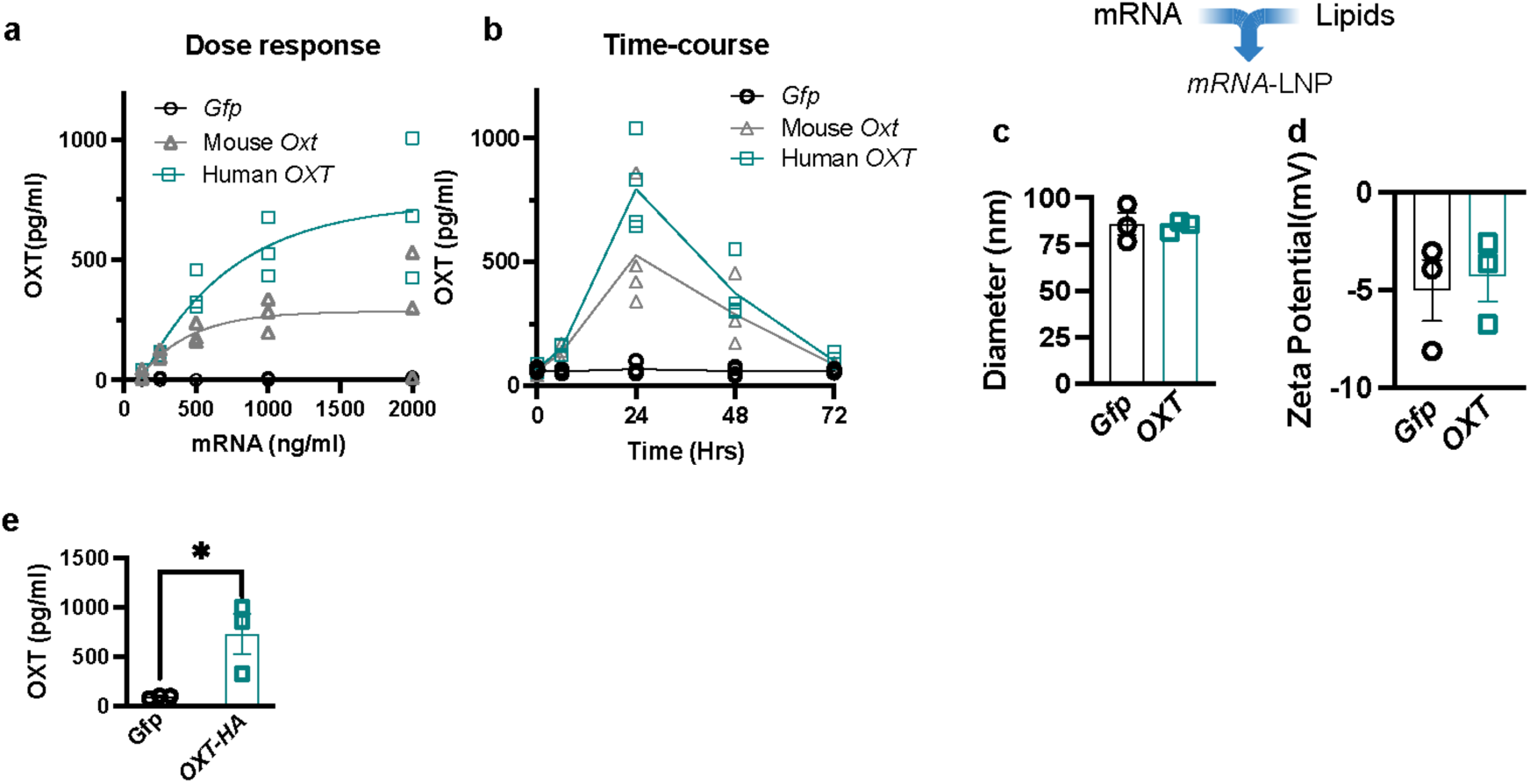
*In vitro* characterisation of mRNA-LNP. **a.** Dose-response curve showing that higher concentrations of oxytocin mRNA led to higher oxytocin peptide concentrations in an *in vitro* setting, with maximal production saturating at 1,000 ng/ml (n=3 per concentration). **b**. Time-course evaluation showing peak oxytocin peptide concentration at 24 hours post-transfection with concentrations tapering off by 72 hours (n=3 per time point). **c-d**. Zetasizer analysis showing LNP size (∼80-100 nm) and zeta potential (∼-2 to −10 mV). **e.** *OXT-HA* transfection in HEK293 cells produces secreted peptide in media, as detected via ELISA. Data presented as mean ± s.e.m. Significance was determined by unpaired T-test comparisons (*p<0.05). n=3 per mRNA construct.

**Fig S4.**
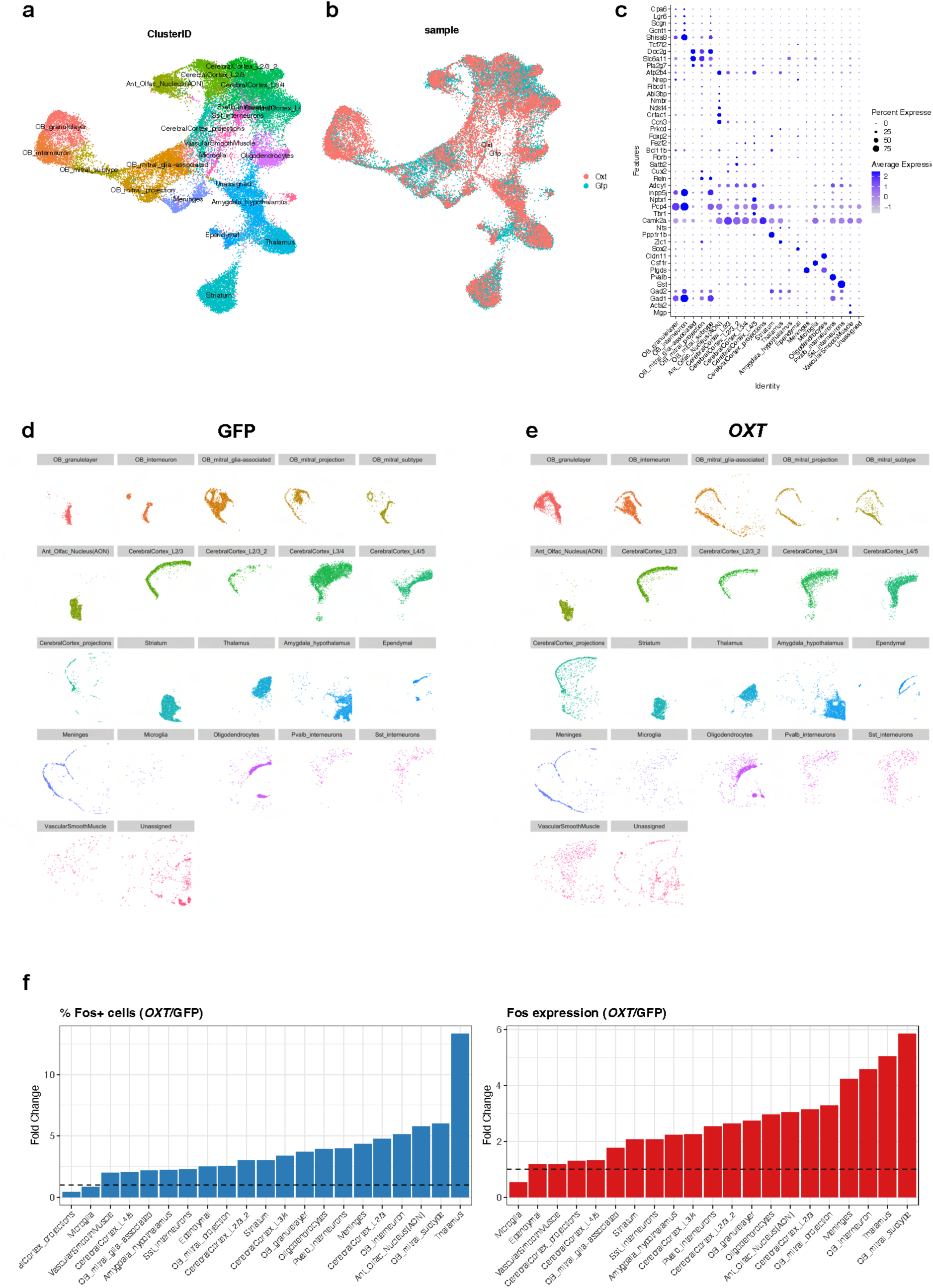
Visium HD Spatial Transcriptomics Analysis. **a.** UMAP projections of 22 clusters that were identified and annotated following analysis with Seurat. Clusters include granule layer, interneuron, mitral glia, mitral projection of the OB, Anterior Olfactory Nucleus (AON), Layer 2-5 and projections of the cerebral cortex, striatum, thalamus, amygdala and hypothalamus, ependymal, meninges, microglia, oligodendrocytes, Pvalb+ and Sst+ interneurons, vascular smooth muscle and a small group of unsassigned cells. **b.** Distribution of cells by treatment condition on UMAP, showing minimal differences. **c.** Marker gene dotplots for each cluster. Spatial distribution of each cluster in *Gfp* (**d**) and *OXT* (**e**) treated brains. **f.** Quantification of % change in *Fos* expressing cells for each cluster. **g.** Quantification of change in normalised *Fos* expression per cluster.

**Fig S5.**
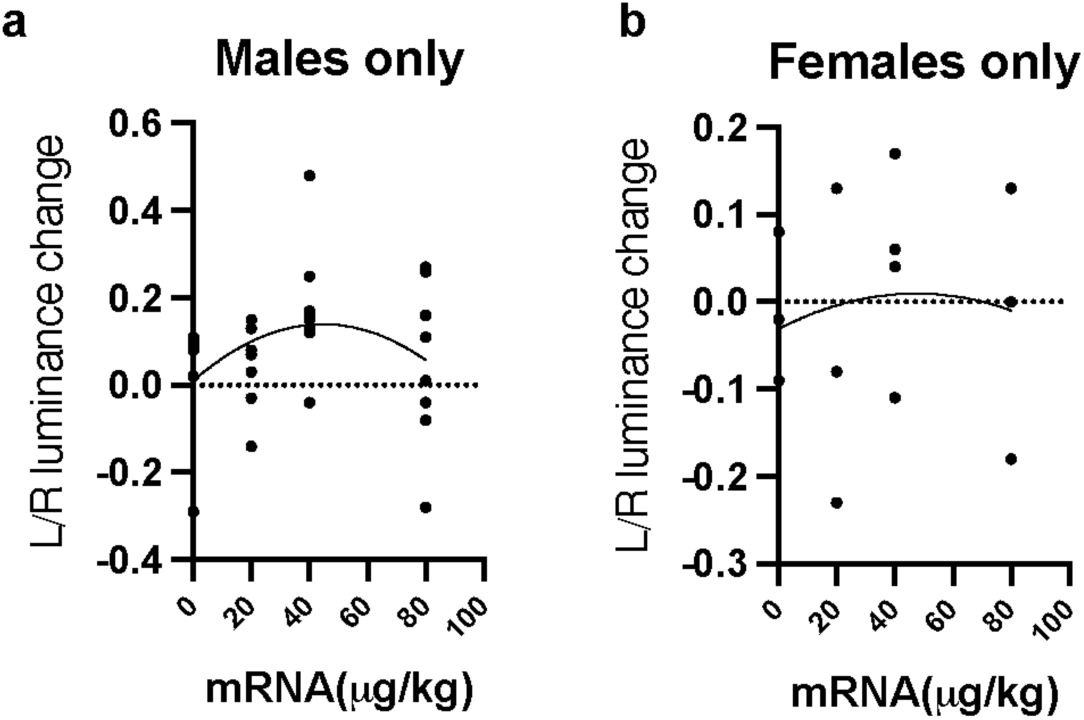
*OXT* mRNA-LNP is analgesic in neuropathic males but not females. **a**. U-shaped dose response in weight-bearing changes observed in male mice with SNI, as determined by Blackbox luminance, but not in females (**b**).

**Table S1.**
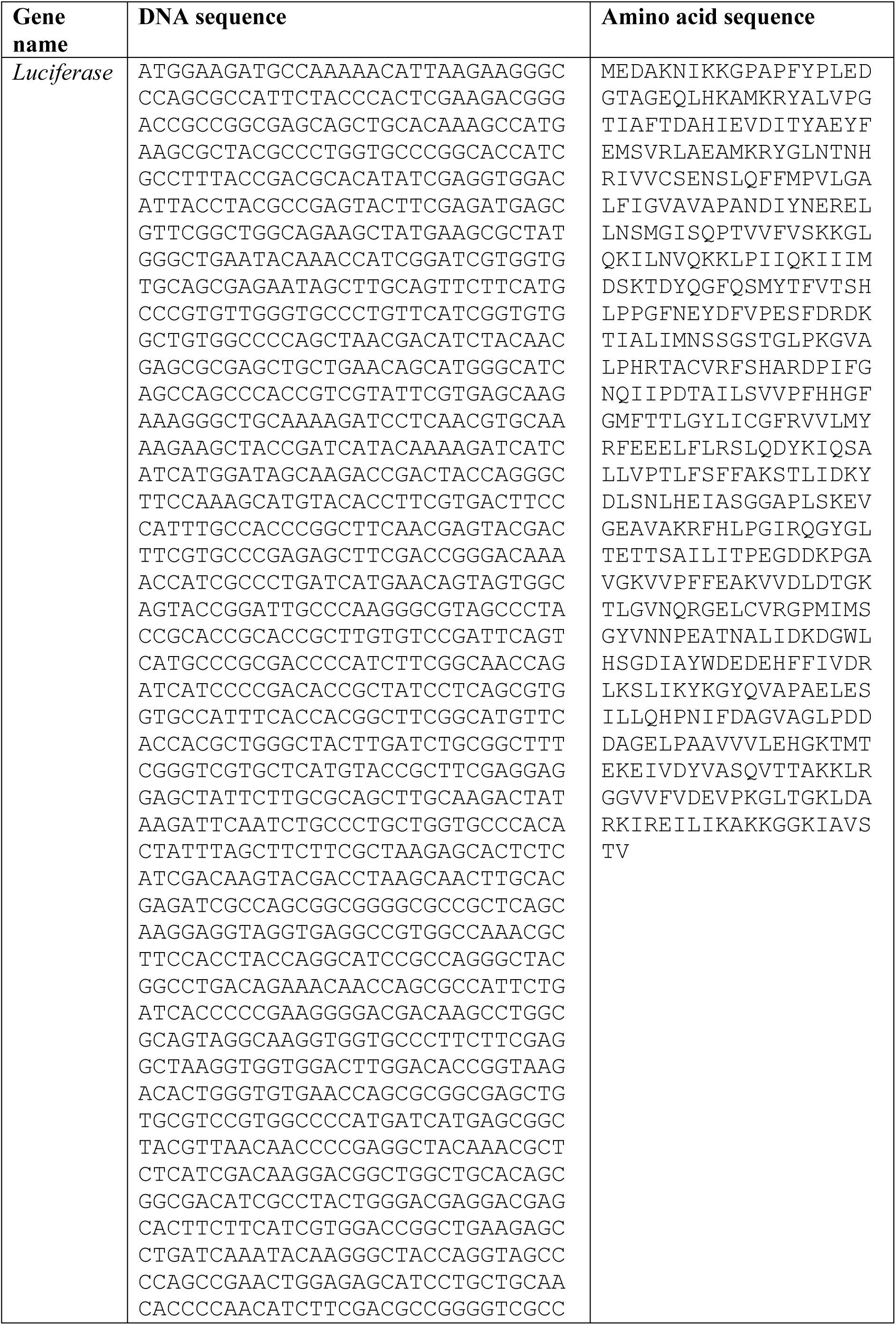
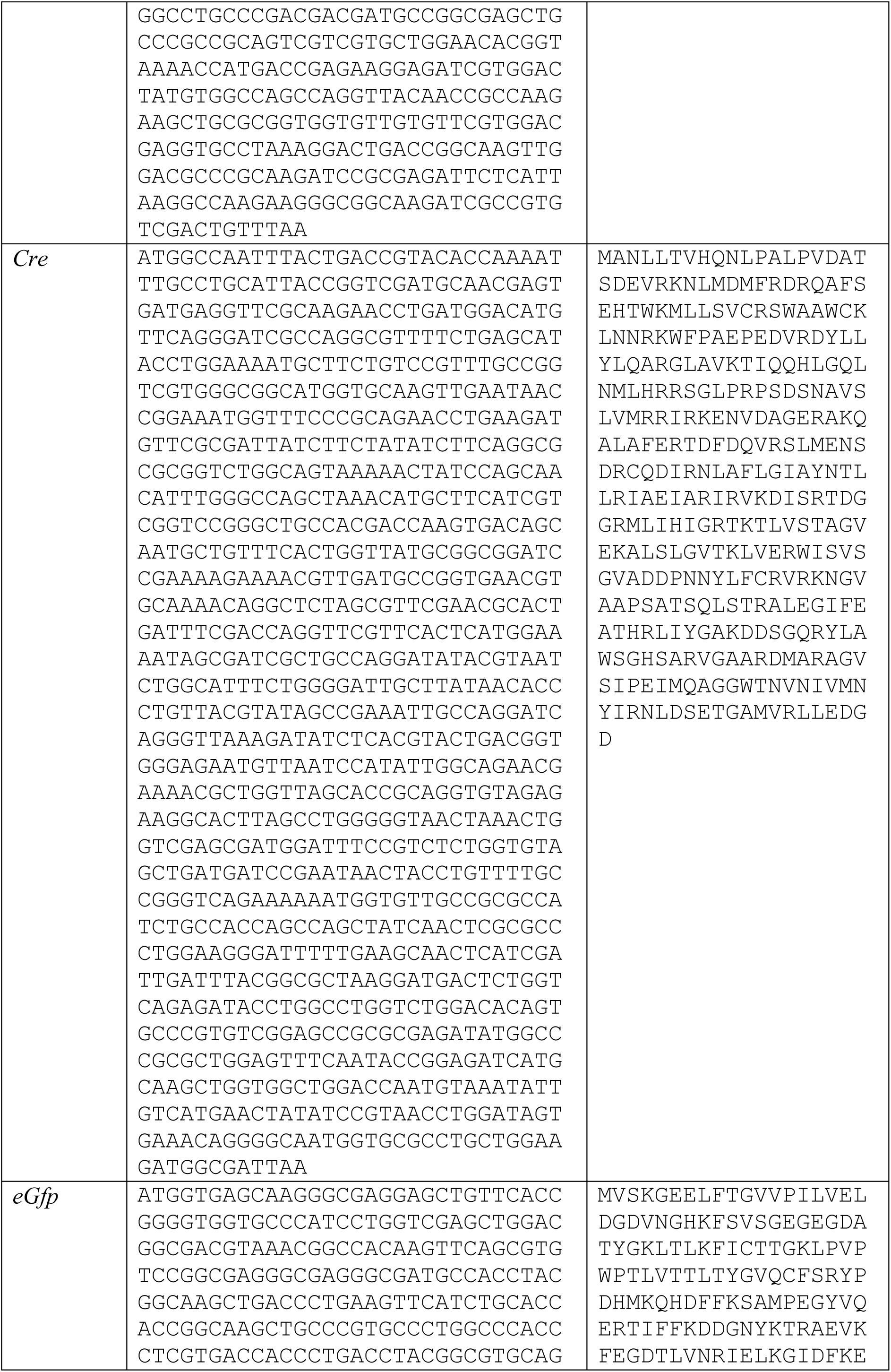
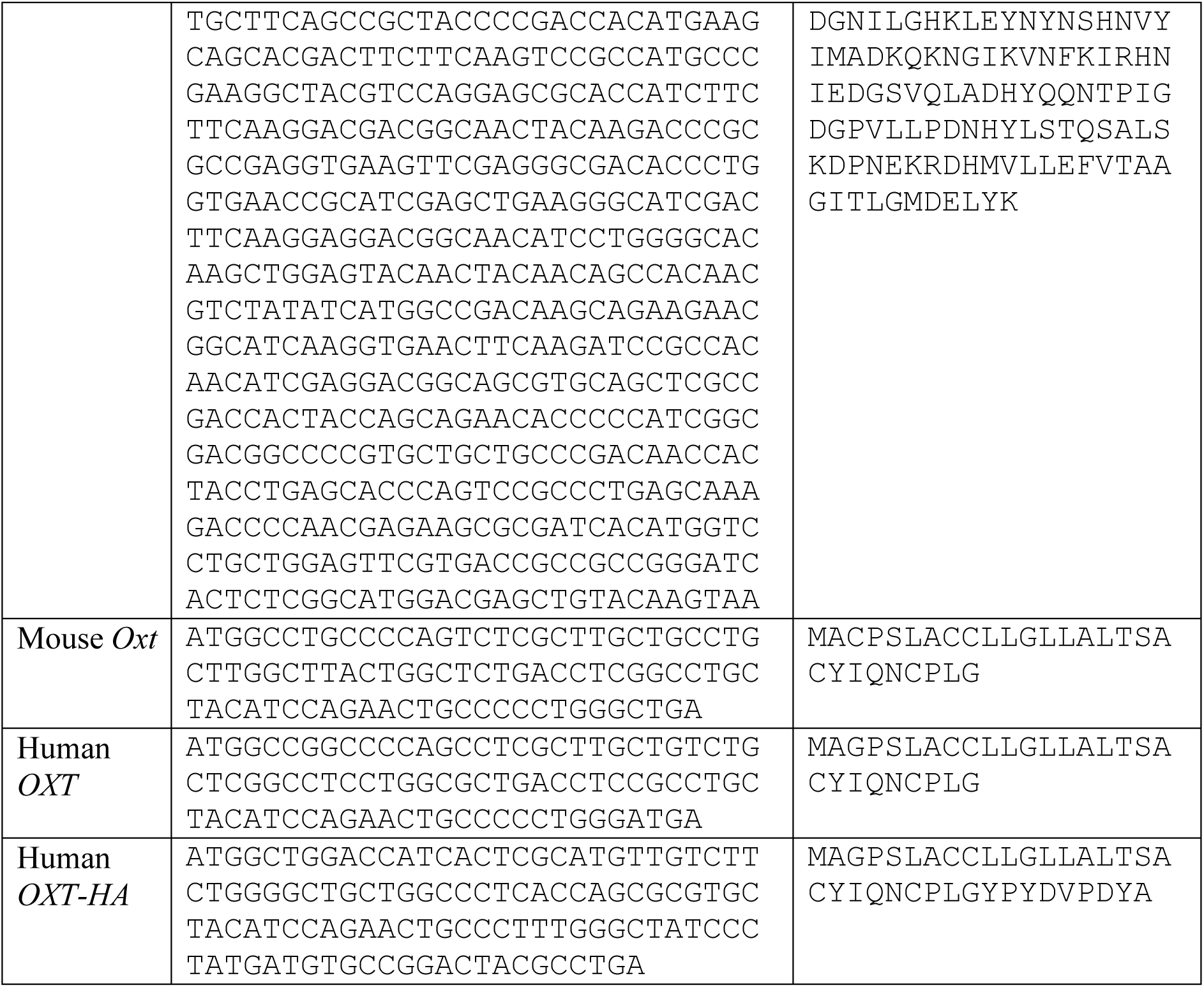
Template sequences and their corresponding protein sequences.

**Table S2.**
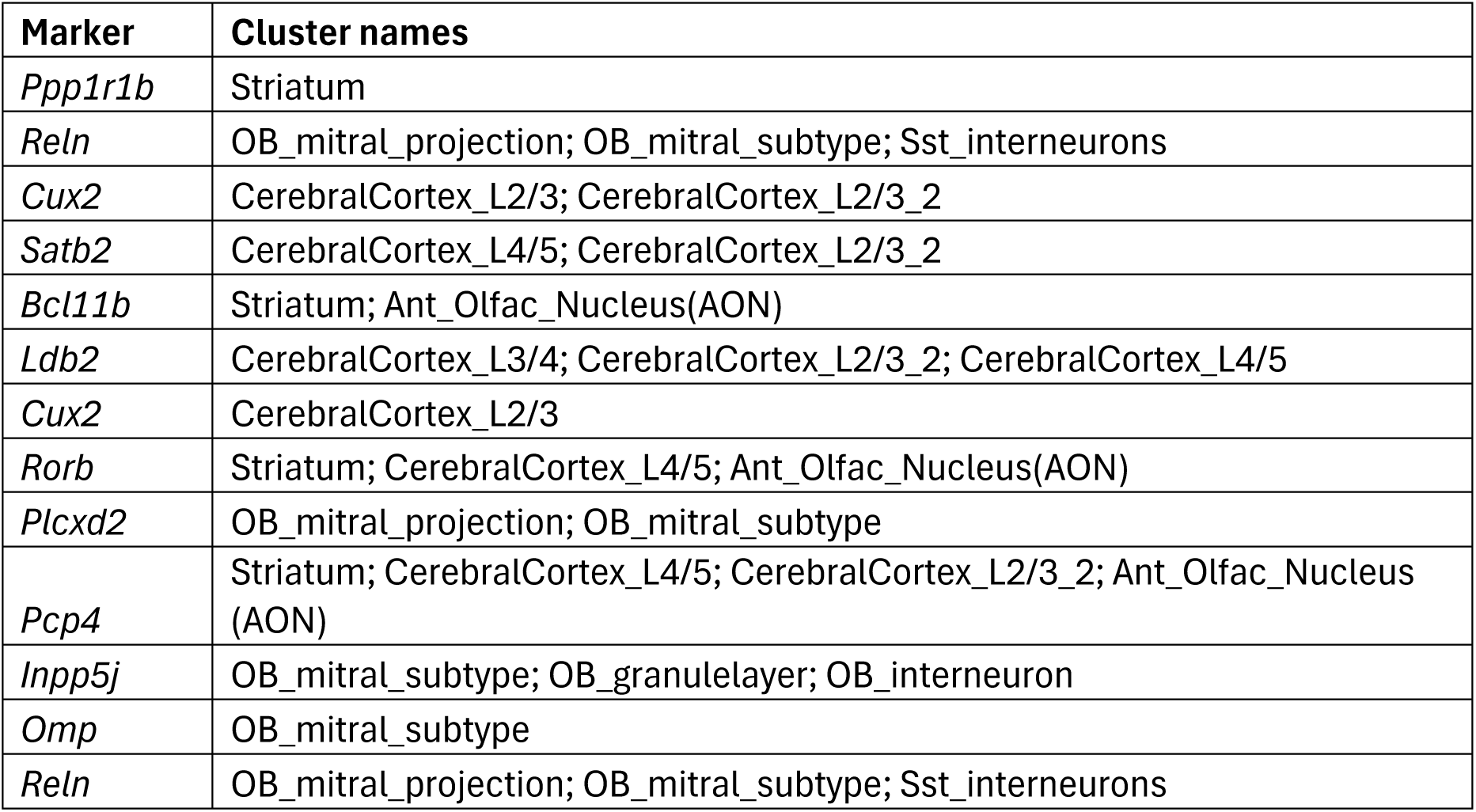
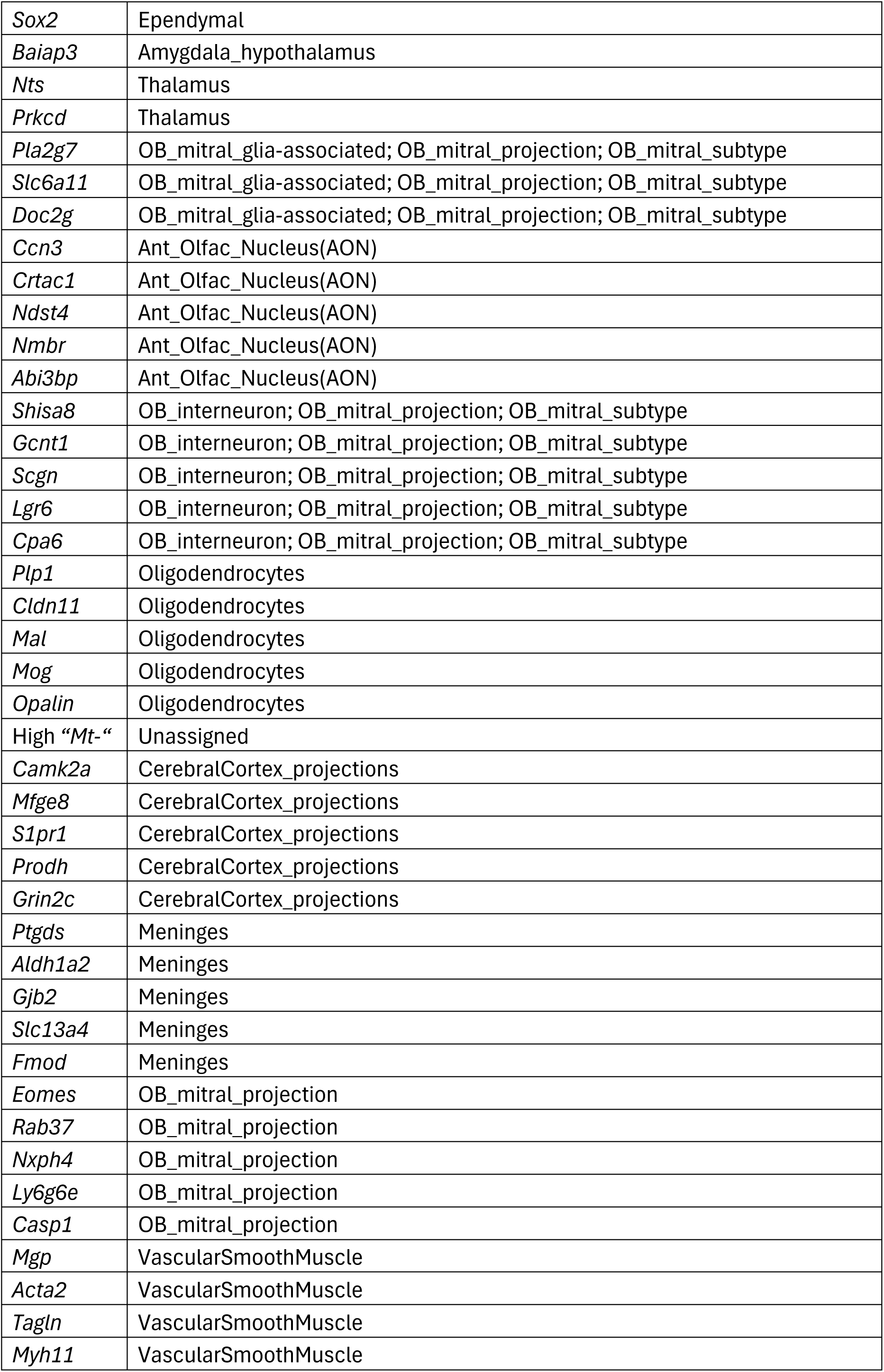
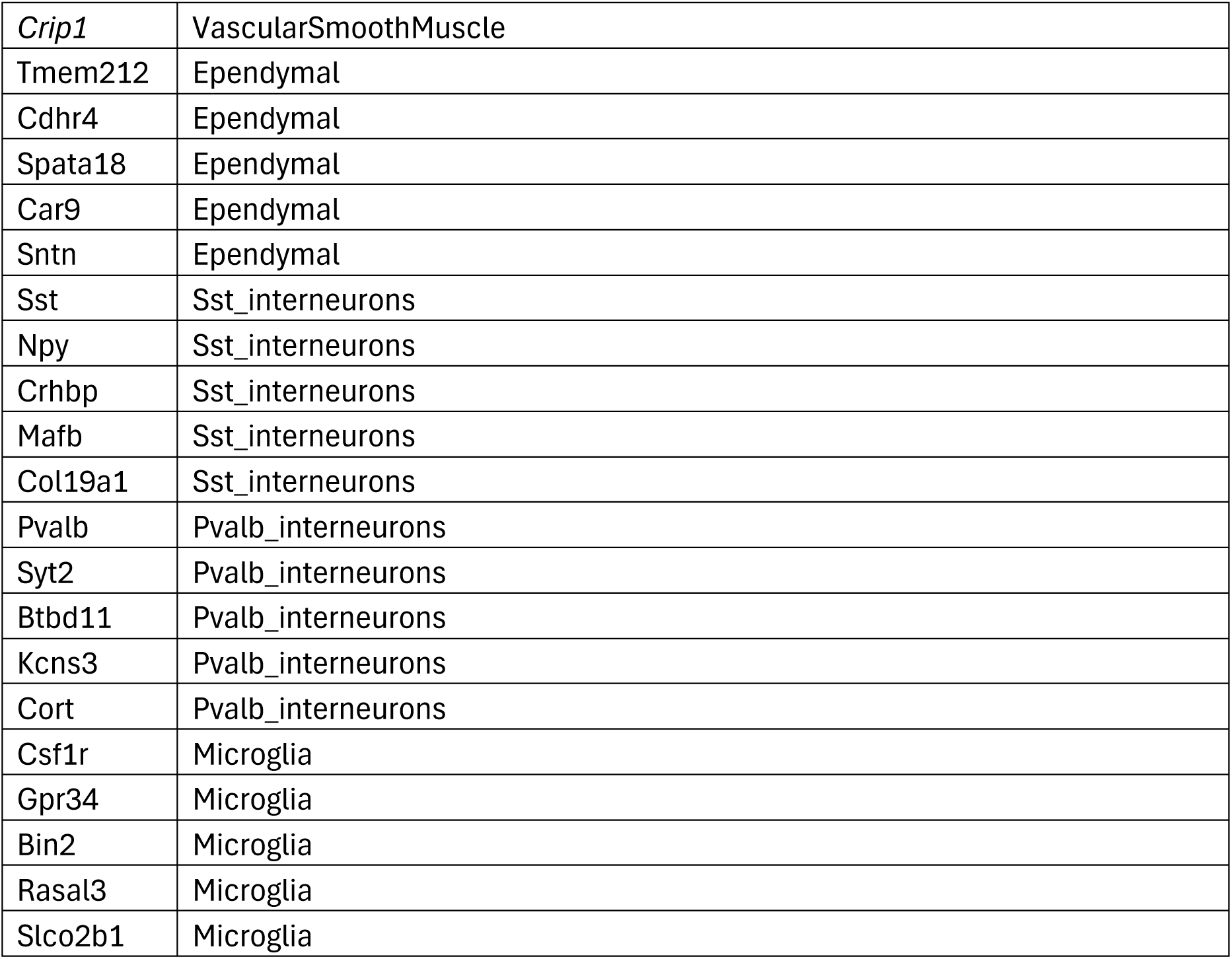
Marker genes used for brain cluster annotation (Visium HD). Summarised in dot plot in Fig S4c.

